# EpiSmokEr: A robust classifier to determine smoking status from DNA methylation data

**DOI:** 10.1101/487975

**Authors:** Sailalitha Bollepalli, Tellervo Korhonen, Jaakko Kaprio, Miina Ollikainen, Simon Anders

## Abstract

**Motivation:** Smoking strongly influences DNA methylation, with current, former and never smokers exhibiting different methylation profiles. To advance the practical applicability of the smoking-associated methylation signals, we used machine learning methodology to train a classifier for smoking status prediction.

**Results:** We show the prediction performance of our classifier on three independent whole-blood test datasets demonstrating its robustness and global applicability. Furthermore, we examine reasons for biologically meaningful misclassifications through comprehensive phenotypic evaluation. The major contribution of our classifier is its global applicability without a need for users to determine a threshold value applicable for each dataset to predict the smoking status.

**Availability and Implementation:** We provide an R package, *EpiSmokEr*, facilitating the use of our classifier to predict smoking status in future studies. *EpiSmokEr* is available from GitHub: https://github.com/sailalithabollepalli/EpiSmokEr

## 1 Introduction

Various studies have assessed the effect of smoking on DNA methylation using microarrays (Ambatipudi, et al., 2016; Elliott, et al., 2014; Gao, et al., 2015; Guida, et al., 2015; Joehanes, et al., 2016; Tsaprouni, et al., 2014; Zeilinger, et al., 2013; Zhang, et al., 2016; Zhang, et al., 2014). The well-established impact of smoking on epigenetic and expression patterns necessitates the inclusion of smoking status as a confounder in DNA methylation or gene expression studies. However, information on the study subjects’ smoking habits is not always available, and where it is, it is not necessarily reliable as subjects often under-report their smoking (Benowitz, et al., 2009). Additionally, current smokers also self-report as non-smokers, which is particularly evident when comparing self-reported smoking with measurements of cotinine, a reliable short-term metabolic marker of nicotine exposure (Benowitz, et al., 2009; Hsieh, et al., 2011). Hence, it has been proposed to determine a subjects’ smoking habits from DNA methylation data itself and subsequently use this information as a covariate in the association study of interest to reduce the potential confounding due to smoking (Shenker, et al., 2013).

To this end, Elliot *et al* (2014) have suggested calculating a weighted score referred to as “smoking score” (SSc) by adding up the methylation levels of the 187 CpG sites found to be significantly associated with smoking by Zellinger *et al* (2013), after first multiplying each methylation value by its effect size in Zellinger et al.’s EWAS study. This ad hoc procedure yields a score that is associated with smoking behavior, yet there is no clear way to find a suitable threshold value to delineate smoking categories across all datasets. Hence, Elliot *et al*. (2014) used different threshold values for two different cohorts on which they tested the score. The subsequent work by Zhang *et al* (2016) used a regression approach. They first determined significantly associated CpG sites in their cohort, and then employed stepwise logistic regression with forward selection. They used a significance-based stopping criterion and found that already after including the 4 most significant loci, the addition of the fifth locus no longer resulted in a significant reduction of the fit deviance. Therefore, their proposed “methylation score” (MS) only requires four CpG sites.

Comparing to standard practices in machine learning (see Hastie *et al* (2009) for a standard textbook) suggests two possible improvements here: First, as the aim is prediction rather than establishing association/causality, feature selection should be done using cross validation rather than significance testing. Second, there is no need to a priori restrict the set of features to select from to those which are significant in marginal (i.e., locus-by-locus) tests (see below). We, therefore, investigated whether we could also gain further improvements to the existing DNA methylation-based smoker identification by employing a commonly used state-of-the-art machine learning approach, namely penalized regression with cross validation, specifically by applying L1-penalized regression (commonly called LASSO; least absolute shrinkage and selection operator) (Tibshirani, 1996) with nested cross validation. The epigenetic clock (Horvath, 2013) serves as a successful example of the use of such penalized regression-based machine learning in the field of epigenetics.

We use LASSO in the classification setting (also known as multinomial or logistic regression): Each subject in the training dataset has one of three class labels -- never smoker, former smoker, or current smoker --, and we seek to find a linear combination of methylation values of selected probes that give (via a logistic link function) a probability value for each of the labels for a test subject. The aforementioned previous works focused only on CpG sites that have been shown to be significantly associated with smoking in a prior GWAS-style analysis. There is, however, no need for this in a machine learning setting; in fact, one typically finds that features that, taken on their own, are not statistically significant, can still contribute to improving the classifier. Instead of assuring statistical significance by marginal (i.e., feature by feature) testing, one uses cross-validation to avoid overfitting. This means that the penalty parameter, which controls the size of the selected feature set, is determined by testing a range of candidate values by repeatedly training models on a subset of the data and testing them on a held-out set (see Methods for details). In contrast, the previous work (Zhang, et al., 2016) has used forward stepwise regression with an arbitrary significance threshold as a stopping criterion. As CV directly estimates the classifier’s predictive performance, it is preferable over a significance-based criterion for deciding on the size of the feature set. This also circumvents the often discussed (see, e.g., (Tibshirani, 1996)) issues of lack of type-I error control inherent to stepwise regression.

Another difference in our method compared to the two previous approaches lies in the decision rule: The SSc (Elliott, et al., 2014) needs to be compared to a specific threshold value to call a person a smoker or a non-smoker, and it is unclear how one should determine this threshold. On the contrary, an appealing feature of logistic regression is that it yields probabilities for the class labels, and the class with the greatest probability will be the classifier’s output, thus circumventing the need for an explicit threshold. Alternatively, the probabilities can be transformed to logarithmic odds, with log odds above zero indicating a probability above 0.5. Hence, multinomial regression implicitly sets the threshold to zero.

We developed *EpiSmokEr*, an easy-to-use R package to facilitate the prediction of smoking status. Given methylation data from microarray assays, it provides predicted smoking status as an output. *EpiSmokEr* can be run not only on whole data sets, but also on isolated individual samples. To be comprehensive, we also provide functions for computing SSc and MS.

## 2 Methods

### 2.1 Datasets

We used the DILGOM dataset (n=514; Supplementary Table 2) to train (Inouye, et al., 2010) and five datasets to test our classifier: the Finnish Twin Cohort (FTC n=408) (Kaprio, 2013), the EIRA study (n=687) (Liu, et al., 2013), the CARDIOGENICS consortium (n=464) (Tsaprouni, et al., 2014) (Supplementary Table 3), and publicly available buccal (Prasad, et al., 2016) and PBMCs datasets (Dogan, et al., 2014) (Supplementary Table 4). For the detailed description of the datasets please refer to Supplementary Methods.

### 2.2 Pre-processing and normalization

All statistical analyses were performed in R. To ensure inclusion of high-quality samples and probes, stringent quality control measures were implemented following the pipeline by Lehne *et al* (2015). To quantile normalize the training dataset, we separated the probe intensity values into six categories based on the color channel, probe-type and subtype. Please refer to the Supplementary Methods for details.

Following normalization, cross-reactive probes (n=27363) from Chen *et al* (2013) and probes with variance < 0.002 (n=372434) across all the individuals were removed. The final training dataset consisted of 52421 probes from 474 samples.

The test datasets were normalized using the same six sets of quantiles (corresponding to probe categories) obtained from the training dataset using a custom function. This approach ensures cross-study performance by fitting the distribution of the test dataset to the training dataset and can be called a “frozen” quantile normalization, following McCall *et al* (2010).

### 2.3 Multinomial LASSO regression

For training the predictor, we used the set of 52,421 high quality probes available from the DILGOM data. We fitted a LASSO-penalized generalized linear model of the multinomial family using the R package glmnet (Friedman, et al., 2010), for which we briefly review the mathematics here. We write *i* for the index of the CpG probe on the array and *j* for the subject (sample). Our input data are the quantile normalized methylation values *x*_*ij*_ (the “beta values” mentioned above) and the subjects’ smoking status categories (class labels) *k*_*j*_ ∈*K* = { never-smoker, former-smoker, current-smoker }. In multinomial regression, the model fits a linear predictor *η*_*jk*_ which is understood as a multinomial equivalent to the log odds of logistic regression, i.e., the predictor corresponds to the probability *p*_*jk*_ which the classifier assigns to the call that subject *j* has smoking status *k* via the logistic transformation,

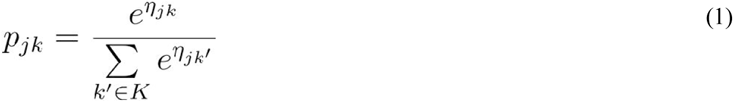

The denominator ensures that the probabilities for the three smoking statuses add up to 1.

The linear predictor η_*jk*_ is given as a linear combination of the fitted model coefficients *β*_*ik*,_

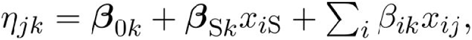

where each probe *i* has three coefficients *β*_*ik*_. The methylation value on the 0–1 scale (often called “beta value”) for probe *i* in sample *j* is denoted *x*_*ij*_. The three intercepts *β*_0*k*_ effectively take up the role of thresholds. As we have included sex as an additional covariate, a sex coefficient *β*_S*k*_ appears in the model which is added to the intercept if subject *i* is male (*x*_*i*S_ = 1) and omitted if subject *i* is female (*x*_*i*S_ = 0).

The coefficients are found by the model fitting procedure, which maximizes the penalized log likelihood,

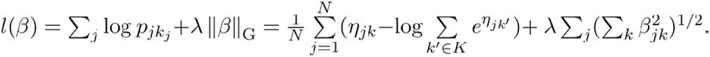

The penalty term, *λ* ‖*β*‖_G_, serves to prevent overfitting. It causes the fit to select only those features (probes) *i* that contribute most to the prediction and shrinks the coefficients *β*_*ik*_ for all other probes to zero. We are using here the “grouped” shrinkage option, which the *glmnet* function offers for multinomial regression. It links the three coefficients for a given probe *i* such that they are selected or not selected together (which is reflected in the penalty term by the penalty acting on 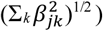

The optimal value for the penalty parameter *λ*, which regulates the strength of this selection force, is chosen by cross-validation, which we describe next.

### 2.4 Cross-validation

An optimal lambda of 0.055 was chosen from a sequence of 100 candidate values by performing 100 iterations of internal cross-validation. In each iteration, the original training dataset was randomly subdivided into 90% training data and 10% hold-out data. A multinomial LASSO regression model was fit on the training dataset, using the candidate values for *λ*. For each lambda value, the fit of the prediction is used on the hold-out data, to obtain estimated probabilities per each class of smoking status.

Specifically: In each iteration, for every value of *λ*, the multinomial deviance

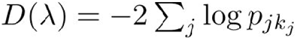

was computed, where the sum runs over the held-out subjects and 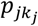 is the probability that the classifier has assigned (using Equation (1)) to held-out subject *j* for that subject’s true smoking status *k*_*j*_ when using the coefficients obtained from training with the given penalty parameter λ. We then averaged the deviances for each λ value over the 100 iterations and chose the λ value with lowest average deviation as optimal λ.

Using this λ, we then performed the final fit on the full training dataset and obtained non-zero coefficients for 121 probes. These coefficients (for each of the three statuses: 121 CpG coefficients, one sex and one intercept coefficients) are used in the calculation of smoking statuses for the test data sets (Supplementary Table 1).

### 2.5 Smoking status prediction

When we use the classifier on a given test data set, that data set first had to be quantile normalized. As stated above, this is done reusing the quantiles obtained in the training set. Then, for each subject in the test data set, a probability is calculated for each of the three smoking statuses using Equation (1). The smoking status category with the highest probability is reported as the classifier’s call.

### 2.6 Secondary analyses

To comprehensively scrutinize the misclassifications of our classifier we used the well-annotated FTC dataset. We examined the duration of smoking abstinence (years since quitting) and cumulative smoking exposure (pack-years), the two most informative smoking behavior variables. Also, we examined the effects of passive smoking on the misclassification.

## 3 Results

### 3.1 Training data

Our training dataset (the DILGOM survey) comprised 474 individuals representative of the Finnish adult population, with a broad age distribution and extensive smoking behavior information (Supplementary Table 2 and Supplementary Methods). We used peripheral blood leukocyte DNA methylation data, assessed by Infinium HumanMethylation450 Bead-Chips, and self-reported smoking status information.

An optimal penalization parameter was determined through internal cross-validation. This resulted in a classifier using 121 CpG sites corresponding to 92 genes (Supplementary Table 5). Figure 1 illustrates the workflow of the classifier with respect to the training and test datasets.

**Fig. 1.**
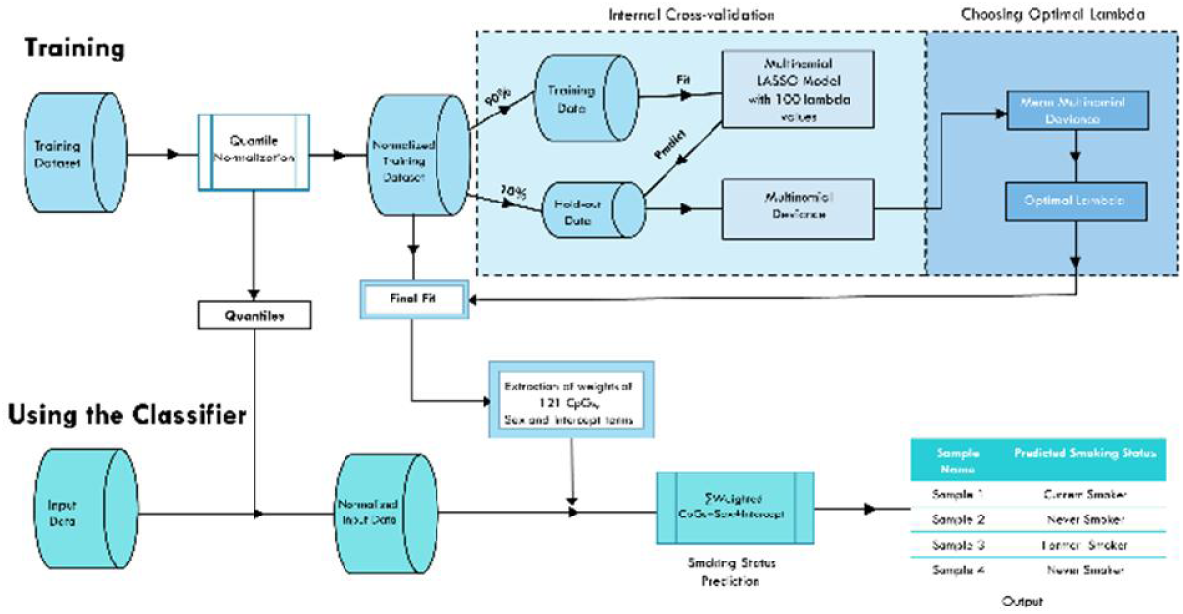
Schematic representation of the workflow of the classifier. *Training*: Quantile normalized training data was used to perform 100 iterations of internal cross-validation to choose the optimal lambda. Using that lambda, a final fit was performed on the full training data, yielding 123 non-zero coefficients. *Using the Classifier*: The input methylation data is first normalized and then probabilities for each smoking status are calculated. The smoking status with the highest probability is taken as the call.

We used sex as an additional covariate owing to differences in smoking prevalence among men and women. We did not adjust for blood cell type composition as SVD analysis revealed no strong association between top principal components and blood cell subtypes (Supplementary Methods). Moreover, several studies have reported that adjusting for blood cell type composition does not alter the smoking-associated methylation signals (Ambatipudi, et al., 2016; Elliott, et al., 2014; Joehanes, et al., 2016; Shenker, et al., 2013; Tsaprouni, et al., 2014).

### 3.2 Validation and performance comparison

For validation, we have chosen three external, independent test datasets: FTC, EIRA and CARDIOGENICS, originating from different populations to demonstrate transferability and global applicability of our classifier (Supplementary Table 3). To visualize the performance of our classifier we generated confusion matrices by comparing the self-reported with predicted smoking behavior (Supplementary Figure 1 A-C). To quantify the performance of our classifier, we determined sensitivity and specificity (Supplementary Table 6). Owing to the multinomial model with three smoking status categories, these values were calculated for each category by comparing that one versus the union of the other two categories (See Methods). On average in the three test datasets, self-reported current smokers were identified with 81% sensitivity and 85% specificity, while never smokers were identified with 94% sensitivity and 57% specificity (Supplementary Table 6). Sensitivity values were highest in the “never vs other” comparison across all three test datasets. For former smokers, the classifier showed low sensitivity across the test datasets, with an average sensitivity of only 18%. However, the classifier was able to identify individuals who did not belong to the former smoker category with a high average specificity of 96%.

As the papers on SSc and MS did not provide a fixed threshold value to use, we tested multiple threshold values for SSc and MS to compare with our method. For SSc a threshold value of zero showed good sensitivity for discriminating current smokers from other smoking status categories (Supplementary Figure 2). For MS a threshold of −7.5 seemed appropriate across all the test datasets to discriminate current from never smokers with an average sensitivity of 85% and specificity of 68% (Supplementary Figure 3). However, it is was not possible to find a single threshold for discriminating former from never smokers with MS that yields reasonable sensitivity and specificity values for all test datasets.

We could not perform extensive comparison of the performance from all the three methods owing to the underlying differences in the approaches (multinomial vs binomial). Our motivation was to predict smoking status, while the other two methods focus on computing a score. To provide a meaningful comparison we calculated sensitivity and specificity values for SSc and MS by first identifying optimal threshold values applicable to test datasets as just described. Our classifier performed well in discriminating current and never smokers from others and showed moderate to marginal ability in discriminating former smokers from others across the test datasets. MS showed good performance across all the test datasets, if used only for a binary classification comparison of discriminating current from never smokers.

To illustrate the results from SSc and MS we plotted the scores against self-reported smoking status (Supplementary Figures 4A and 4B). We also show that the threshold of 17.55 proposed by SSc (Elliott, et al., 2014) for European ethnicity failed to distinguish the majority of the current smokers from other smoking status categories (Supplementary Figure 4 A). While (Elliott, et al., 2014) suggested that different ethnicities might require different thresholds, this was not the case for our classifier. For instance, the training dataset used for building the classifier was from the Finnish population and yet the classifier performed well in the test datasets originating from different populations.

Results from our classifier are also shown as ternary plots to visualize the smoking probabilities predicted by our classifier for all three classes simultaneously (Supplementary Figures 4C-4E). Each corner of a ternary plot corresponds to a smoking status category. Each point in the ternary plot represents an individual with a corresponding triplet of estimated smoking probabilities which add up to 100%. So, the higher the probability for a specific smoking status category the closer the point is to the corresponding corner. Individuals with mixed profiles, showing agreement with more than one category, are on one of the side lines or in the central part of the ternary plot. Hence, we observe most of the former smokers in the middle of the ternary plot as they exhibit mixed profiles of current and never smokers.

### 3.3 Reasons for misclassification

To understand the possible reasons for misclassification we performed a comprehensive phenotypic assessment on the two test datasets with additional phenotypic information. In FTC and EIRA, the “current smoker” category has been further subdivided into current daily smokers and occasional smokers. To highlight the impact of occasional smokers on the performance of the classifier, we have also shown the confusion matrices with the occasional smokers as a separate category in the FTC and EIRA datasets (Supplementary Figure 1 D-E). For instance, in the EIRA dataset, of the 66 occasional smokers, 53 were predicted as never smokers and 6 as former smokers (Supplementary Figure 1E). This misclassification can be attributed to the similar methylation profiles of occasional smokers to that of never and former smokers. Therefore, we expect a decrease in the classification accuracy when current daily and occasional smokers are combined into current smoker category.

We also observed an increase in the sensitivity values when occasional smokers were excluded from the current smokers. For example, the exclusion of 66 occasional smokers in the EIRA dataset improved the sensitivity from 69% to 88% (Supplementary Table 8). Similarly, in the FTC dataset exclusion of 6 occasional smokers increased the sensitivity from 82% to 87%. To also understand the misclassification of former smokers, we used the extensive smoking behavior information available from the FTC dataset. In FTC, the former smokers class had the highest misclassification rate, with 72% (n=101) of former smokers identified as never smokers (Supplementary Figure 1D and Supplementary Figure 5 A-C). Phenotypic verification revealed that 85 of 101 individuals had quit smoking more than 10 years prior to blood sampling for DNA methylation measurement (Supplementary Figure 5 A-B). The 6% (n=9) of former smokers identified as current smokers had quit smoking recently and had higher mean pack-years compared to other former smokers (Supplementary Figure 5 B-C). We also observed lower mean and range for pack-years among current smokers identified as never smokers (Supplementary Figure 5B).

In addition to active smoking, every class of smokers were exposed to passive (second-hand) smoking to a certain extent (Supplementary Figure 5D). We also noticed that some of the individuals were exposed to both active and passive smoking for prolonged periods (Supplementary Figure 5D). Our classifier results indicated that methylation levels reflect the cumulative smoking exposure. However, from our analyses, it is hard to determine the extent of passive smoking which leads to a change in the methylation levels.

### 3.4 Performance in buccal tissue and PBMCs

We have used methylation data from buccal tissue (Prasad, et al., 2016) and peripheral blood mononuclear cells (PBMCs) (Dogan, et al., 2014) to assess the performance of the classifier. Current smokers were identified with the highest sensitivity (95%) and specificity (97%) in the buccal tissue dataset (Supplementary Table 4). Based on the self-reported smoking status, this dataset used the labels “cigarette smoker” and “non-tobacco smoker”. Specificity drops by 37% when considering samples that were identified as former smokers by our classifier. This suggests that around 38% (n=15) of the non-tobacco smokers are former smokers based on their methylation profiles. This is in line with the specific cohort definition of non-tobacco smokers (Prasad, et al., 2016) being individuals abstinent from tobacco or nicotine-containing products for at least 5 years.

Similarly, in the PBMC dataset with 50 smokers and 61 non-smokers, sensitivity is reduced to 76% if we were to consider the inclusion of samples that were classified as former smokers by our classifier. Notably, both the buccal tissue and PBMC datasets comprised individuals of African-American ethnicity, demonstrating the global applicability of our classifier across different ethnicities.

### 3.5 The R package *EpiSmokEr*

We implemented the SSt classifier as an R package. To estimate smoking behavior, *EpiSmokEr* (*Epi*genetic *Smok*ing status *E*stimato*r*) expects methylation data from the Infinium HumanMethylation450 beadchip arrays as input. Input data can be either raw methylation data in the form of intensity data (IDAT) files or a normalized methylation matrix on the beta scale ranging from 0 to 1. A sample sheet with sex is required to complement the methylation data. The *normaliseData* function has a suite of customized internal functions to normalize and calculate beta values from the IDAT files. To perform SQN and ILN normalization we use functionality from the minfi package (Aryee, et al., 2014), and for QN we use a custom function. Quantiles from our training dataset are used to adjust the distribution of the input data. Our classifier estimates the probabilities of an individual being a current, former or never smoker. The output from the classifier is a label, namely the smoking status category with the highest probability. *EpiSmokEr* has been already tested on several datasets containing 400 to 700 samples and takes only a few minutes starting with the IDAT files to the estimation of smoking status. An additional advantage of *EpiSmoker* is that it offers functions for computing smoking and methylation scores, providing users with a choice for their analyses. *EpiSmokEr* is available from GitHub: https://github.com/sailalithabollepalli/EpiSmokEr

## 4 Discussion

To address the need for predicting smoking status reliably, we built a DNA methylation based classifier using a machine learning approach. Our SSt classifier calculates probabilities of an individual being a current, former or never smoker. The smoking status category with the highest probability is reported as the predicted smoking status. A major advantage of our approach is its general applicability without a need for an ad hoc threshold as a decision boundary.

All the three methods compared here share the common objective of translating methylation at smoking-responsive CpGs to a measure reflecting smoking behavior. The SSc (Elliott, et al., 2014) focuses on distinguishing self-reported current smokers from other classes while the MS (Zhang, et al., 2016) can only distinguish current from never smokers and former from never smokers. Our SSt approach focuses on determining smoking status considering all the three smoking categories. Moreover, users do not need to determine a threshold specific for their dataset and can characterize a single sample to large datasets within few minutes starting with raw methylation files.

### 4.1 Performance evaluation

We evaluated the performance of our classifier by calculating sensitivity and specificity across the test datasets by comparing self-reported smoking status with predicted smoking status. As the calculation of sensitivity and specificity requires a threshold applicable to all datasets and smoking status categories we tried to determine optimal threshold values for SSc and MS. Our classifier showed good discriminative ability for identifying current smokers and never smokers across all the test datasets. MS (Zhang, et al., 2016) performed well across the test datasets, however, this approach can only distinguish between two smoking status categories. Notably, the European ethnic smoking score threshold of 17.55 proposed earlier (Elliott, et al., 2014) was not applicable in any of the datasets used here. This suggests that the threshold is dataset-specific and may be partly governed by technical or genetic differences which cannot be discerned. Also, in the case of SSc and MS, a threshold score needs to be determined each time and for each dataset to distinguish smoking categories, which makes it difficult to interpret the score when calculated for individual samples as there is no reference value to compare with.

In addition to the differences in the training schemes used in the three smoking behavior prediction, it is essential to consider other factors that might have affected their performance. Sensitivity and specificity measures from our classifier are lower than those reported by SSc (Elliott, et al., 2014). However, these methylation biomarkers identified for smoking were mainly focused on classifying heavy smoking (Elliott, et al., 2014). To make a fair comparison we must consider factors such as cohort composition, gender, age, and the smoking status considered by the other classifiers. For deriving the European ethnic SSc threshold of 17.55 to discriminate current smokers from others (Elliott, et al., 2014), 16 heavy smokers from a dataset of 95 men were considered. To develop the MS(Zhang, et al., 2016) older adults aged 50–75 years with long smoking history were used. On the other hand, we have trained our classifier on a dataset with a broad age spectrum, including both men and women. MS (Zhang, et al., 2016) considers only binary comparisons, while our SSt focuses on identifying the smoking behavior of an individual giving equal priority to all the three smoking classes.

The performance metrics may also be affected by the fact that we calculate the prediction accuracy estimates by comparing the predicted status to self-reported smoking behavior, which is prone to misreporting and poor recall of long-term smoking history (Benowitz, et al., 2009). Furthermore, the approaches are based on the independence (mutually exclusive) assumption that each smoking status category has a unique methylation profile without any overlap with other categories. However, this assumption may not be valid for smoking-associated methylation profiles and also for smoking behavior in general.

DNA methylation profiles of former smokers might resemble either current or never smokers. Results from the secondary analyses in the FTC dataset using total time of abstinence and cumulative pack-years showed that misclassifications of the former smokers by our classifier are dependent on how long they have been abstinent and how much and for how long they have smoked before quitting. Misclassification of former smokers with longer cessation time as never smokers has also been reported earlier (Ambatipudi, et al., 2016; Elliott, et al., 2014; Guida, et al., 2015; Joehanes, et al., 2016; Tsaprouni, et al., 2014; Zeilinger, et al., 2013). This specific misclassification can be attributed to the reversal of methylation levels in former smokers to the levels similar to never smokers as a function of cessation time.

Also, current smokers can greatly vary based on their current and cumulative past smoking behavior, as well as the use of tobacco products other than cigarettes. We observed that current occasional smokers are often misclassified as never smokers. Their classification clearly depended on when they have last smoked, how much they typically smoke on days that they smoke and the frequency of smoking. This was also evident in the FTC and EIRA datasets, where the classifier performance was negatively affected by including occasional smokers.

Apart from misreporting and passive smoking, the never smoker category is expected to be more homogenous compared to the other two categories. Passive smoking was prevalent across all the smoking status categories in the FTC dataset. However, based on the results from our classifier, we cannot quantify the extent of the impact of passive smoking on the methylation profiles and prediction of smoking status.

The heterogeneity within each category leads to mixed DNA methylation profiles, which is biologically and perhaps clinically relevant, and as expected. However, the same may lead to misclassification, which could be falsely interpreted as poor performance of the classifier.

To sum up, it is important to consider the biological relevance and the aforementioned factors while interpreting the results and performance metrics from a classifier. Although our classifier considers three smoking status categories, the intrinsic limitations associated with former smokers and DNA methylation was also reflected in our results. Varying definitions of former smokers across different datasets might also impact the sensitivity of the classifier. Given the complexity of former smokers both biologically and behaviorally, we caution that former smokers may remain as an ultimate challenge for any DNA methylation-based prediction algorithm. Nevertheless, from a clinical/biological perspective, despite the misclassification, results from the classifier are still significant as they reflect the impact of smoking on DNA methylation and potentially gene function of the individual.

### 4.2 Performance in other tissues

Given the tissue-specific nature of DNA methylation, the good performance of our whole blood data trained classifier on the buccal tissue data was perhaps surprising. However, a recent study (Nwanaji-Enwerem, et al., 2018) implemented the smoking index (SI) based on 66 blood-derived smoking-related CpGs (Gao, et al., 2016) on the same buccal dataset and showed successful discrimination of smokers from non-smokers. Our results are also in line with another study that showed a substantial overlap between smoking-associated CpGs identified in blood and buccal tissues obtained from matched samples (Teschendorff, et al., 2015).

Similarly, results from the PBMCs are also reassuring, as these cells are extracted from whole blood. An EWAS conducted on this dataset (Dogan, et al., 2014) identified highest methylation difference between smokers and non-smokers for two loci in the AHRR gene (cg05575921 and cg23576855), which were also a part of the 121 CpGs of our classifier. Considerable performance of our classifier in buccal tissue and PBMCs suggests a broader and systematic impact of smoking on the methylome. Interestingly the buccal dataset used here comprised samples of Caucasian and African-American origins. Hence, this dataset served as a case study to demonstrate the global applicability of our classifier. However, further testing in large datasets from multiple tissues is required to confirm the cross-tissue performance of our classifier.

### 4.3 Practical implications

Identification of smoking status based on methylation is more robust than other traditional biomarkers with short half-lives in body fluids. Our classifier provides an objective measure of smoking status and is globally applicable curtailing the need for computing population/ethnic/study-specific thresholds. In addition to the predicted smoking status, results from the classifier include probabilities belonging to each smoking category. For former smokers, these probabilities can also be used to understand possible misclassification.

Predicted smoking status from our classifier can be useful when self-reported data is missing or not available or is presumed to be highly inaccurate or to validate self-reported data. This is advantageous for larger consortia studies and biobanks where a smoking phenotype was not collected at the time of sample collection. Methylation-derived smoking status can also serve as a tool in forensic settings and can also be used as a covariate to adjust for smoking-associated confounding in association analyses like EWAS and GWAS.

## 5 Conclusions

We have developed a robust DNA methylation-based predictor providing an objective measure of smoking status. Our approach considers three smoking status categories and is applicable to datasets from different populations. We also examined reasons for misclassifications through comprehensive phenotypic evaluation. We provide the community with an R package, *EpiSmokEr*, where raw or normalized DNA methylation data can be used to predict smoking status. *EpiSmokEr* includes functionality to also calculate SSc and MS allowing users to select the most appropriate method and output for their purposes.

## Acknowledgements

We thank all the researchers for making their datasets publicly available. We are grateful to the DILGOM and the FTC for providing the data. We thank all the participants who contributed to different cohorts. We warmly acknowledge Dr Milla Kibble and Dr Emma Cazaly for providing constructive feedback on the manuscript.

---

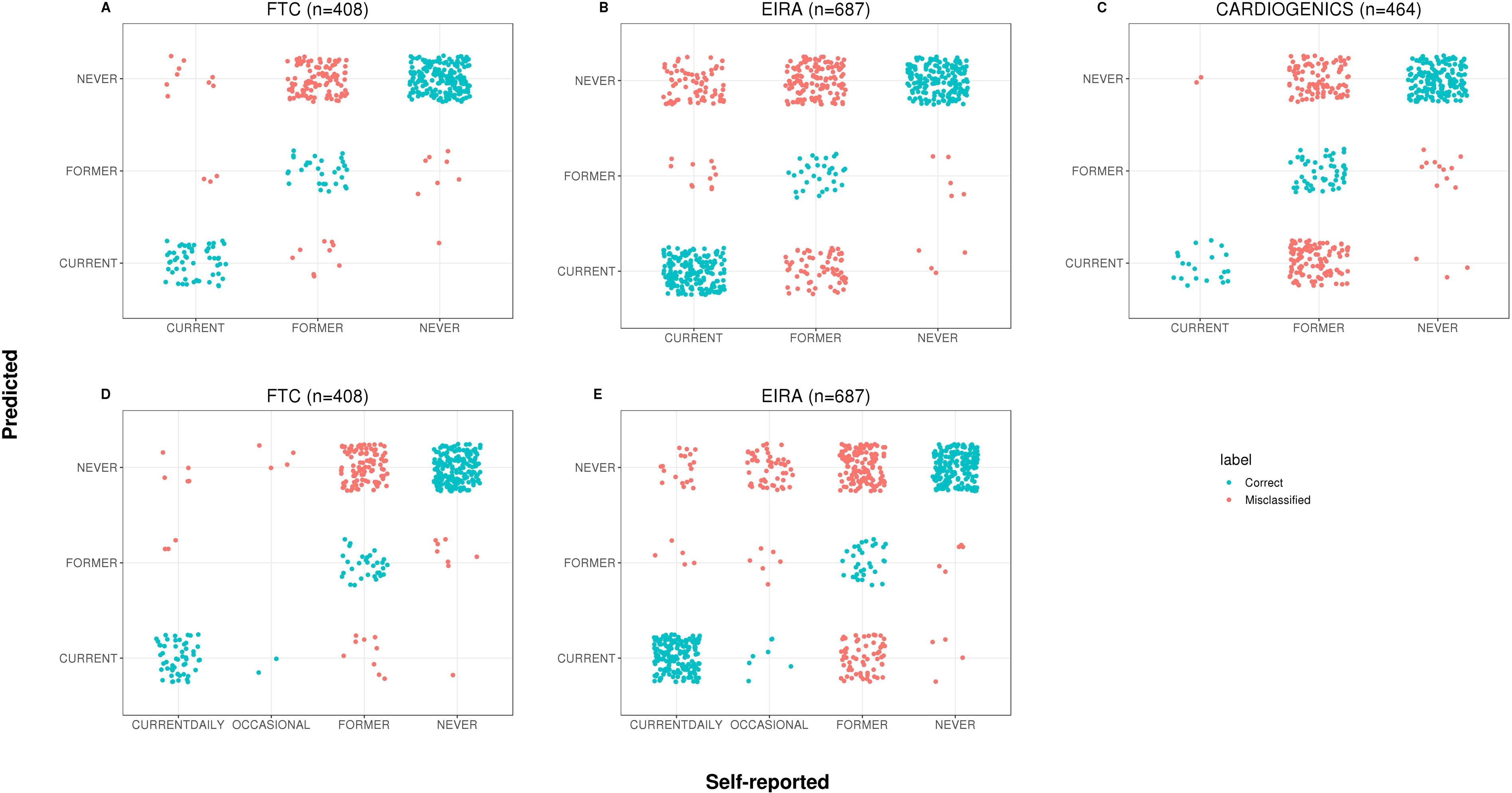

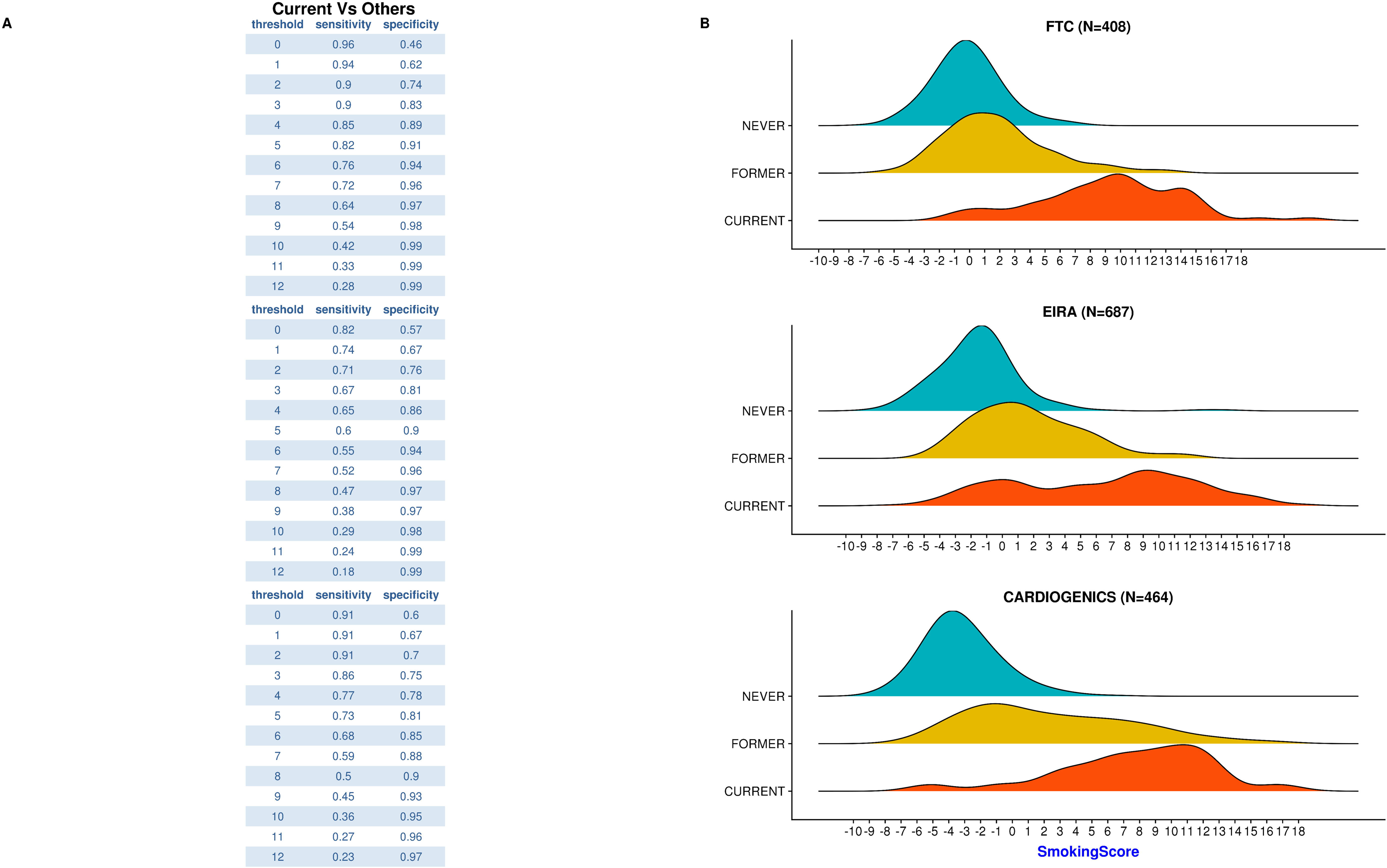

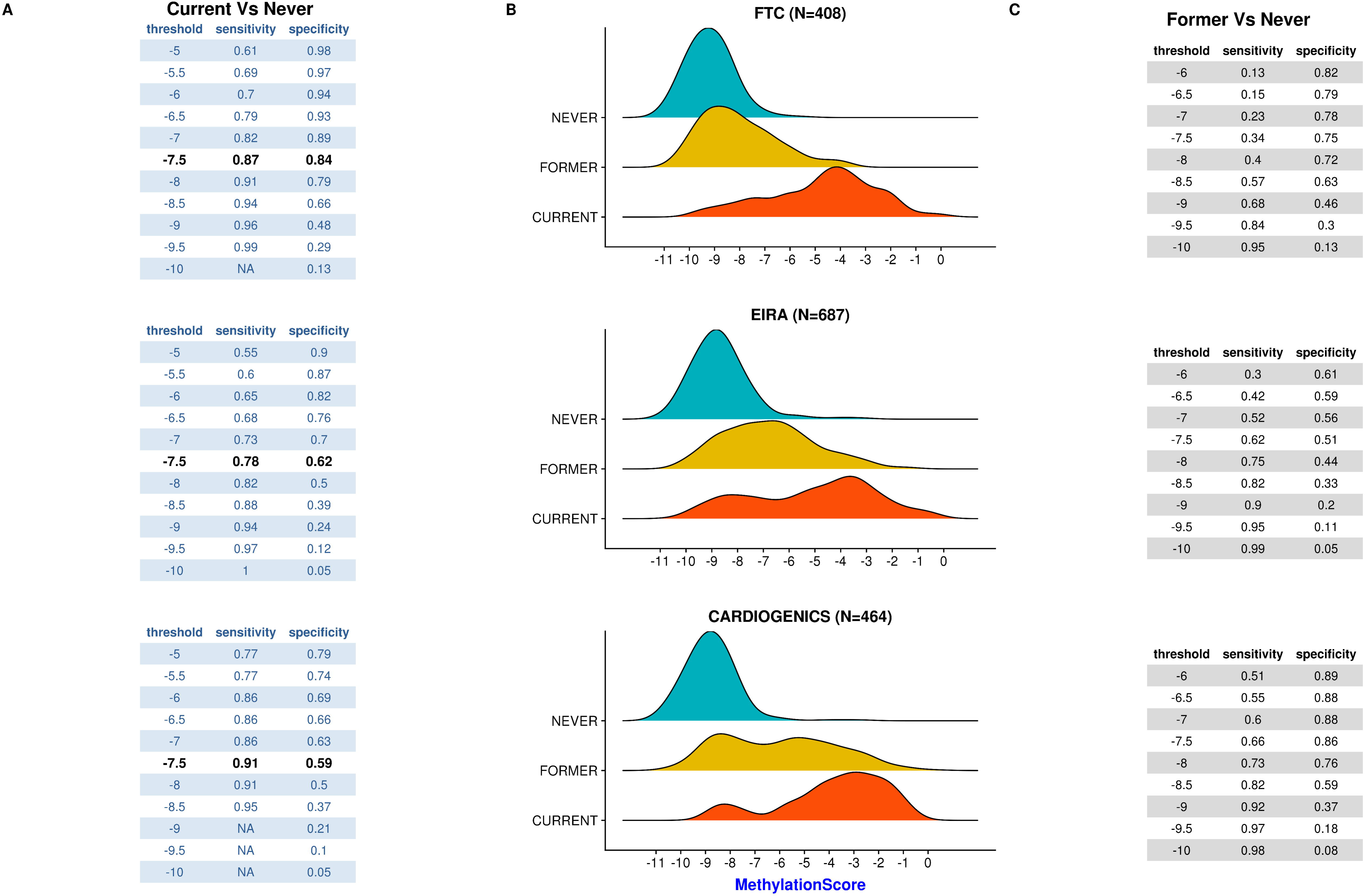

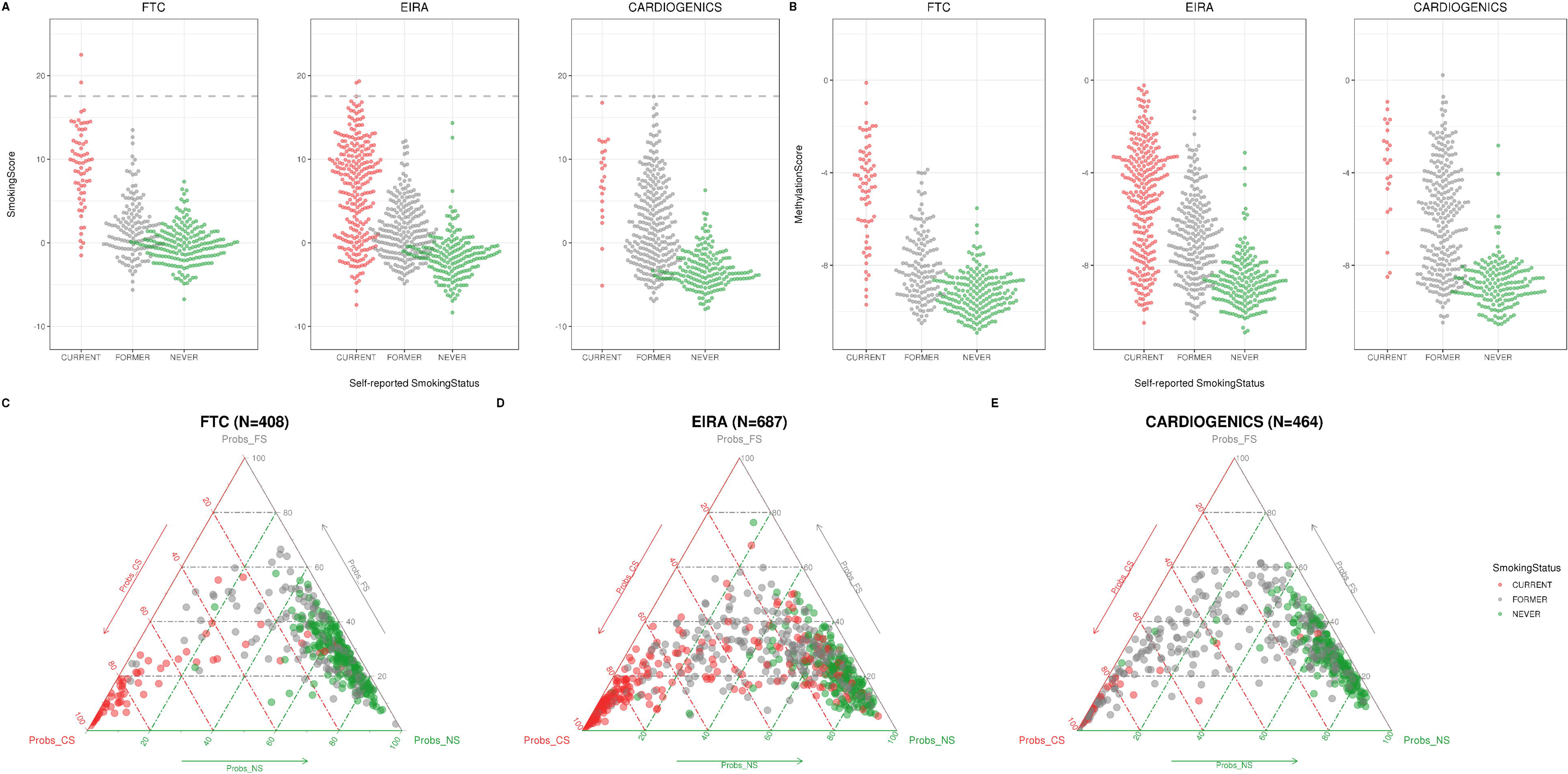

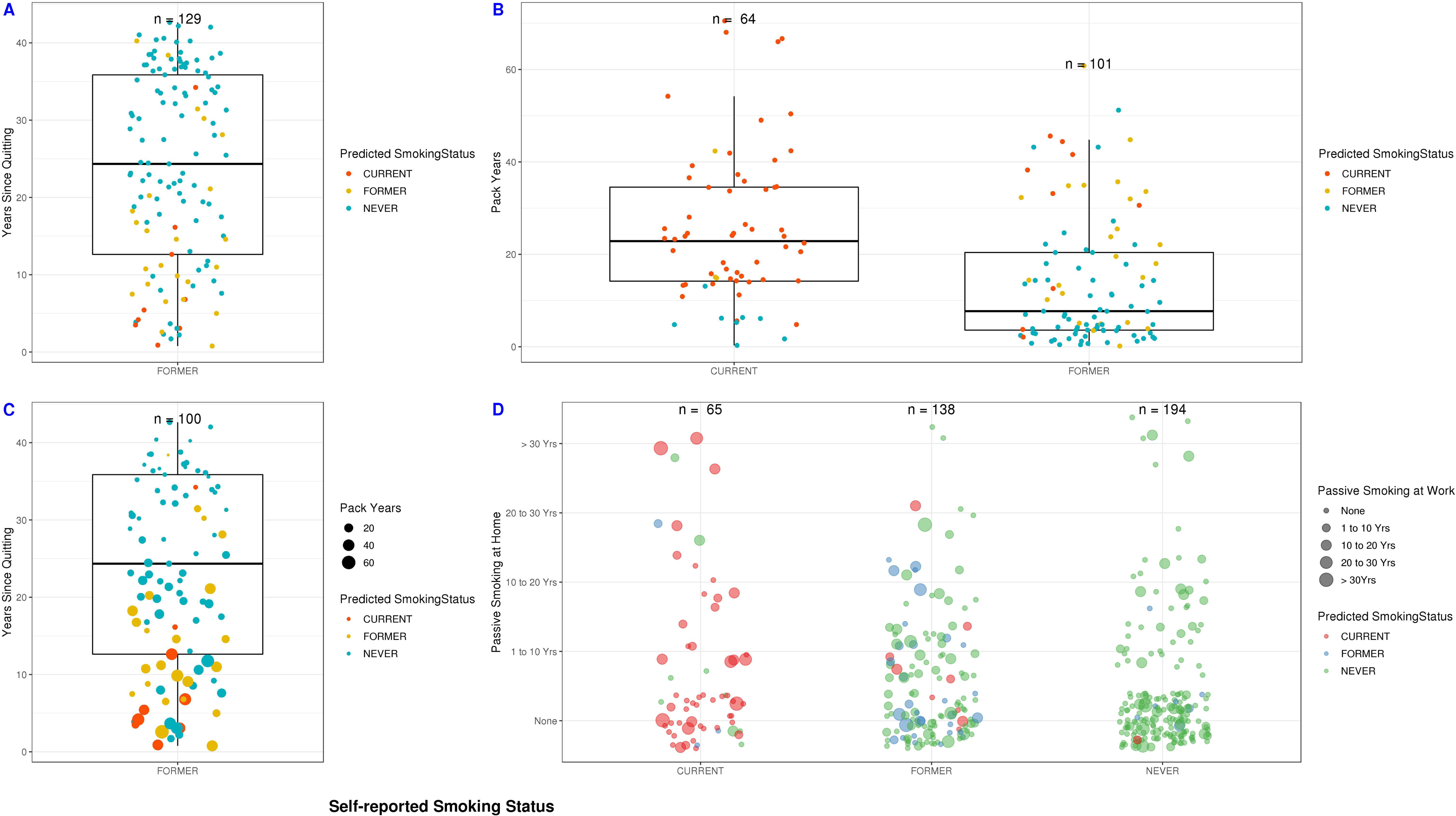

## References

Ambatipudi, S., et al. (2016) Tobacco smoking-associated genome-wide DNA methylation changes in the EPIC study, Epigenomics, 8, 599–618.

Aryee, M.J., et al. (2014) Minfi: a flexible and comprehensive Bioconductor package for the analysis of Infinium DNA methylation microarrays, Bioinformatics, 30, 1363–1369.

Benowitz, N.L., et al. (2009) Prevalence of Smoking Assessed Biochemically in an Urban Public Hospital: A Rationale for Routine Cotinine Screening, Am. J. Epidemiol., 170, 885–891.

Chen, Y.-A., et al. (2013) Discovery of cross-reactive probes and polymorphic CpGs in the Illumina Infinium HumanMethylation450 microarray, Epigenetics, 8, 203–209.

Dogan, M.V., et al. (2014) The effect of smoking on DNA methylation of peripheral blood mononuclear cells from African American women, BMC Genomics, 15, 151.

Elliott, H.R., et al. (2014) Differences in smoking associated DNA methylation patterns in South Asians and Europeans, Clin. Epigenetics, 6, 4.

Friedman, J., Hastie, T. and Tibshirani, R. (2010) Regularization Paths for Generalized Linear Models via Coordinate Descent, J. Stat. Softw., 33, 1–22.

Gao, X., et al. (2015) DNA methylation changes of whole blood cells in response to active smoking exposure in adults: a systematic review of DNA methylation studies, Clin. Epigenetics, 7, 113.

Gao, X., et al. (2016) Relationship of tobacco smoking and smoking-related DNA methylation with epigenetic age acceleration, Oncotarget, 7, 46878–46889.

Guida, F., et al. (2015) Dynamics of smoking-induced genome-wide methylation changes with time since smoking cessation, Hum. Mol. Genet., 24, 2349–2359.

Hastie, T., Tibshirani, R. and Friedman, J. (2009) The Elements of Statistical Learning: Data Mining, Inference, and Prediction, Second Edition. Springer Science & Business Media.

Horvath, S. (2013) DNA methylation age of human tissues and cell types, Genome Biol., 14, R115.

Hsieh, S.J., et al. (2011) Biomarkers increase detection of active smoking and secondhand smoke exposure in critically ill patients, Crit. Care Med., 39, 40–45.

Inouye, M., et al. (2010) Metabonomic, transcriptomic, and genomic variation of a population cohort, Mol. Syst. Biol., 6, 441.

Joehanes, R., et al. (2016) Epigenetic Signatures of Cigarette Smoking, Circ. Cardiovasc. Genet., 9, 436–447.

Kaprio, J. (2013) The Finnish Twin Cohort Study: an update, Twin Res. Hum. Genet., 16, 157–162.

Lehne, B., et al. (2015) A coherent approach for analysis of the Illumina HumanMethylation450 BeadChip improves data quality and performance in epigenome-wide association studies, Genome Biol., 16, 37.

Liu, Y., et al. (2013) Epigenome-wide association data implicate DNA methylation as an intermediary of genetic risk in rheumatoid arthritis, Nat. Biotechnol., 31, 142–147.

McCall, M.N., Bolstad, B.M. and Irizarry, R.A. (2010) Frozen robust multiarray analysis (fRMA), Biostatistics, 11, 242–253.

Nwanaji-Enwerem, J.C., et al. (2018) Relationships of Long-term Smoking and Moist Snuff Consumption with a DNA Methylation Age Relevant Smoking Index: An Analysis in Buccal Cells, Nicotine Tob. Res.

Prasad, G.L., et al. (2016) A cross-sectional study of biomarkers of exposure and effect in smokers and moist snuff consumers, Clin. Chem. Lab. Med., 54, 633–642.

Shenker, N.S., et al. (2013) Epigenome-wide association study in the European Prospective Investigation into Cancer and Nutrition (EPIC-Turin) identifies novel genetic loci associated with smoking, Hum. Mol. Genet., 22, 843–851.

Shenker, N.S., et al. (2013) DNA methylation as a long-term biomarker of exposure to tobacco smoke, Epidemiology, 24, 712–716.

Teschendorff, A.E., et al. (2015) Correlation of Smoking-Associated DNA Methylation Changes in Buccal Cells With DNA Methylation Changes in Epithelial Cancer, JAMA Oncol, 1, 476–485.

Tibshirani, R. (1996) Regression Shrinkage and Selection via the Lasso, J. R. Stat. Soc. Series B Stat. Methodol., 58, 267–288.

Tsaprouni, L.G., et al. (2014) Cigarette smoking reduces DNA methylation levels at multiple genomic loci but the effect is partially reversible upon cessation, Epigenetics, 9, 1382–1396.

Zeilinger, S., et al. (2013) Tobacco smoking leads to extensive genome-wide changes in DNA methylation, PLoS One, 8, e63812.

Zhang, Y., et al. (2016) Self-reported smoking, serum cotinine, and blood DNA methylation, Environ. Res., 146, 395–403.

Zhang, Y., et al. (2014) F2RL3 methylation as a biomarker of current and lifetime smoking exposures, Environ. Health Perspect., 122, 131–137.

