## Supplementary Material for "EpiSmokEr: A robust classifier to determine smoking status from DNA methylation data"

### 1 Supplementary Methods

#### 1.1 Datasets and Definition of smoking habits

Below we briefly describe the smoking-status definitions used in the training and test datasets to determine the smoking behavior of participants. Each cohort used self-reported questionnaires to ascertain the smoking habits of the participants. As most of the test datasets we used are from a public repository (GEO), we had limited information on the exact questions that were used in these cohorts to describe smoking behavior.

***Training dataset:*** We used the DILGOM (Dietary, Lifestyle and Genetic determinants of Obesity and Metabolic syndrome) data from the Finnish population-based FINRISK 2007 study as the training dataset (Supplementary Table 2). A detailed description of the DILGOM dataset is provided in greater detail elsewhere (Inouye, et al., 2010; Inouye, et al., 2010). Genome-wide DNA methylation data and self-reported smoking status were available from 514 individuals. Based on the self-reported smoking behavior, three major classes of smoking statuses were identified: current, former and never smokers. In addition, plasma cotinine measures determined by gas chromatography-mass spectrometry (Broms, et al., 2012) were available from 86 current, 31 former and 3 never smokers. For the individuals with cotinine measures available, self-reported smoking status was validated using cotinine levels.

Smoking behavior of the DILGOM study participants was thoroughly assessed using three measures:

1. Self-reported smoking status
a. Never Smoker
b. Former smoker (quit more than a year ago)
c. Recent quitter (quit 1 month - year ago)
d. Current occasional smoker
e. Current daily smoker
2. When did you had your last cigarette? (measured on a scale of 1 to 7)
   1. Yesterday or today
   2. Two days ago to one month ago
   3. One month to half a year ago
   4. Half a year to year
   5. 1 to 5 years ago
   6. 6 to 10 years ago
   7. More than ten years ago
3. Cotinine measurements

A combination of these three measures was used to select individuals for training the classifier.
Self-ascribed current daily smokers and cotinine verified current smokers (with cotinine levels above 10 ng/ml) were considered as current smokers. Current occasional smokers with either high cotinine values or who smoked their last cigarette less than a half a year ago were also considered as current smokers. Former smokers were defined as individuals who quit smoking at least a year ago. Following these criteria, 28 former and 2 never smokers with high cotinine values were excluded from the analyses.

***Whole-blood Test datasets***

We have considered three whole-blood datasets quantified on Infinium HumanMethylation450 BeadChip for validation of our classifier (Supplementary Table 3).

*1. Finnish Twin Cohort (FTC):*

We used 408 twins from the FTC corresponding to the FinnTwin16 study (Kaprio, 2013), a population-based longitudinal study of five consecutive birth cohorts (1975–1979) of Finnish twins and their families. Extensive smoking information including cotinine, cumulative pack-years, duration of smoking abstinence (years since quitting) and passive smoking information was available from these subjects.

Smoking behavior was ascertained on a scale of 1 to 7 as follows:
1 = I smoke 20 or more cigarettes per day
2= I smoke 10 to 19 cigarettes per day
3= I smoke < 10 cigarettes per day
4= I smoke at least once per week but not every day
5= I smoke less than once a week
6= I am in abstinence or have quit smoking
7= I have never smoked

Based on the above scale we classified the participants as follows: 1 to 5 as current, 6 as former and 7 as never smokers. We further classified current smokers as current daily (1 to 3) and occasional smokers (4 to 5).

*2. Epidemiological Investigation of Rheumatoid Arthritis (EIRA):*

EIRA is a population-based case-control study of rheumatoid arthritis conducted in middle and southern parts of Sweden (Stolt, et al., 2003). A total of 687 participants with methylation data, comprising 354 rheumatoid arthritis (RA) cases were used as a test dataset (Liu, et al., 2013).

Smoking behavior of participants was quantified based on the following questions: (1) Do you smoke? (2) If you do not smoke, have you previously smoked? (3) If you have previously smoked, which year did you stop smoking? (4) If you smoke or previously have smoked, when did you start to smoke regularly? (5) How much do you smoke, or did you smoke before you stopped, on average per day?

For RA cases based on the index year (the year in which symptoms of RA onset occurred), current smokers were defined as subjects who smoked during the index year, while ex-smokers (former) were the subjects who stopped regular smoking for at least a year before the index year. Never smokers were the subjects who had never smoked before or during the index year. The controls included were from the same population matched for age, gender and smoking status (Liu, et al., 2013).

*3. CARDIOGENICS:*

We used methylation data available from 464 subjects participating in the CARDIOGENICS Consortium (Tsaprouni, et al., 2014), a European descent case-control study of coronary artery disease (CAD). Ex-smokers (n=263) were defined as individuals who ceased smoking more than 12 weeks (n=251) or less than 12 weeks (n=12) prior to the recruitment day.

Raw intensity files (IDAT) were available for the FTC and EIRA datasets. The CARDIOGENICS dataset was provided in the methylated and unmethylated signal intensity format, suitable only for performing quantile normalization (color channel, probe type and subtype). Self-reported smoking status was available from all of these test datasets.

***Test datasets from tissues other than whole blood***
To investigate and assess the performance of our classifier in other tissues, we used two publicly available normalized datasets from buccal tissue (Prasad, et al., 2016) and PBMCs (Dogan, et al., 2014) (Supplementary Table 4). No normalization step was performed on these datasets owing to the different tissue types.

The buccal tissue data comprised 120 healthy men from North Carolina, comprising 40 long-term smokers, 40 moist snuff users, and 40 nonsmokers (Prasad, et al., 2016). Most of the participants were Caucasians followed by African-Americans (Supplementary Table 4). We have excluded snuff users from our analyses. Following criteria were used in this dataset to define smokers and nonsmokers:
(A) Smokers: individuals who exclusively smoked cigarettes with at least 10 cigarettes/ day for at least 3 years and had an expired carbon monoxide of 10–100 ppm.
(B) Nonsmokers: individuals who reported abstinence from any nicotine-containing or tobacco products for at least 5 years and had an expired carbon monoxide of 0–5 ppm.

The PBMC dataset (Dogan, et al., 2014) consisted of adult African American females from the states of Iowa and Georgia who participated in the Family and Community Health Study (FACHS) (Cutrona, et al., 2005). PBMC samples were collected during the tenth wave of the FACHS. In this dataset, actively smoking individuals were characterized as “smokers” and those denied using any tobacco products were characterized as “non-smokers”.

##### 1.2 DNA methylation analysis

For all whole-blood datasets (Inouye, et al., 2010; Kaprio, 2013; Liu, et al., 2013; Tsaprouni, et al., 2014), bisulfite conversion had been performed using the EZ-96 DNA Methylation-Gold Kit (Zy-mo Research, Irvine, CA, USA) and genome-wide DNA methylation was quantified on the Infinium HumanMethylation450 BeadChip (Illumina, San Diego, CA, USA).

##### 1.3 Pre-processing and Quality control

On the training dataset, we performed quality control following the pipeline by Lehne *et al* (2015). Probes with SNPs, non-CpG probes and probes at sex chromosomes were removed (n=15558). Probes with nominal detection P value above $10^{-16}$ were set to missing. Samples with missing data above 5% and probes with missing data above 2% (n=18509) were excluded. After the QC, four samples were excluded owing to low-quality.

##### 1.4 Quantile normalization

To normalize the training dataset, we separated the probe intensity values into six categories based on the color channel, probe-type and subtype. Within each probe category, quantile normalization was performed by first sorting the actual intensity values within each sample. After sorting, average intensity values were calculated for each rank across all the samples. Then the actual intensity values were replaced with the corresponding averaged intensity values (quantiles), thereby forcing all the samples to have the same distribution. The quantiles obtained from the six probe categories were saved. Beta values were calculated for each CpG probe using normalized intensity values as beta= (Methylated/(Methylated+Unmethylated+100)). We note that we do not rescale the intensities of Infinium II probes on the basis of Infinium I probes, as it is nonessential for our classification purposes.

##### 1.5 Singular Value Decomposition (SVD) Analysis

We performed SVD analysis to check if the top 20 principal components of the DILGOM methylation data were associated with the proportions of blood cell subtypes. The champ.SVD function from R ChAMP package (Morris, et al., 2014) was used to perform this analysis. No strong associations were identified between the PCs and the blood cell types. Only PC-2 had a nominal association with CD8+ T-cells.

##### 1.6 Data preparation

To compare the performance of our classifier with other methods we have normalized the test datasets according to the original publications. For calculating the smoking score (SSc) proposed by Elliott *et al* (2014) we have used subset quantile normalization (SQN) (Touleimat and Tost, 2012). In SQN, first the reference quantiles are computed for each probe category of Infinium I signals based on CpG annotations (S shore, S shelf, N shore, N shelf and distant), to which the type II probes are normalized. To calculate methylation score (MS) proposed by Zhang *et al* (2016), the datasets were normalized using Illumina normalization method (ILN). In ILN, internal controls are used as a reference to which the data were normalized. No additional background correction was performed.

#### 1.7 Smoking scores (SSc) and methylation scores (MS)

SSc are calculated as described by the Elliot *et al.* (Elliott, et al., 2014), which we briefly recapitulate here: for a given subject *j* in a test data set, the smoking score is calculated as

$$\mathrm{SSc}_j = \sum_{i=1}^{187} w_i (x_{ij} - x_i^{\mathrm{NS}})$$

where the index *i* runs over the 187 CpGs listed in Zellinger et al.'s Table S2 (Zeilinger, et al., 2013). As before, we denote with $x_{ij}$ the normalized methylation value of this subject for probe *j*. The reference methylation value, $x_i^{\mathrm{NS}}$, for probe *i*, is Zellinger et al.'s median methylation values of never smokers, averaged over discovery and replication cohorts (columns P and S of their Table S2). The probe weight $w_i$ is given by

$$w_i = \frac{x_i^{\mathrm{CS}} - x_i^{\mathrm{NS}}}{\sum_{i'=1}^{187} (x_{i'}^{\mathrm{CS}} - x_{i'}^{\mathrm{NS}})} \quad ,$$

where $x_i^{\mathrm{CS}}$ is the median methylation values of current smokers, also averaged over discovery and replication cohorts (columns Q and T of their Table S2).

For the MS calculation, the methylation values of 4 CpG probes *cg05575921, cg05951221, cg02451831* and *cg06126421* are multiplied with their corresponding weights given in Figure 4 of Zhang *et al* (2016) and then summed up.

#### 1.8 R packages used

The *preprocessIllumina* and *preprocessQuantile* functions from the minfi package (Aryee, et al., 2014) were used to perform ILN and SQN normalization methods, respectively. The Biobase package (Huber, et al., 2015) was used to save the methylation and phenotype data as an *eSet* object. The glmnet package (Friedman, et al., 2010) was used to perform multinomial LASSO. The following R packages were used to generate figures: ggplot2 (Wickham, 2016), ggtern (Hamilton and Ferry, 2018), ggpubr (Kassambara, 2018), ggbeeswarm (Clarke and Sherrill-Mix, 2017) and cowplot (Wilke, 2018).

#### 2 Supplementary Figures

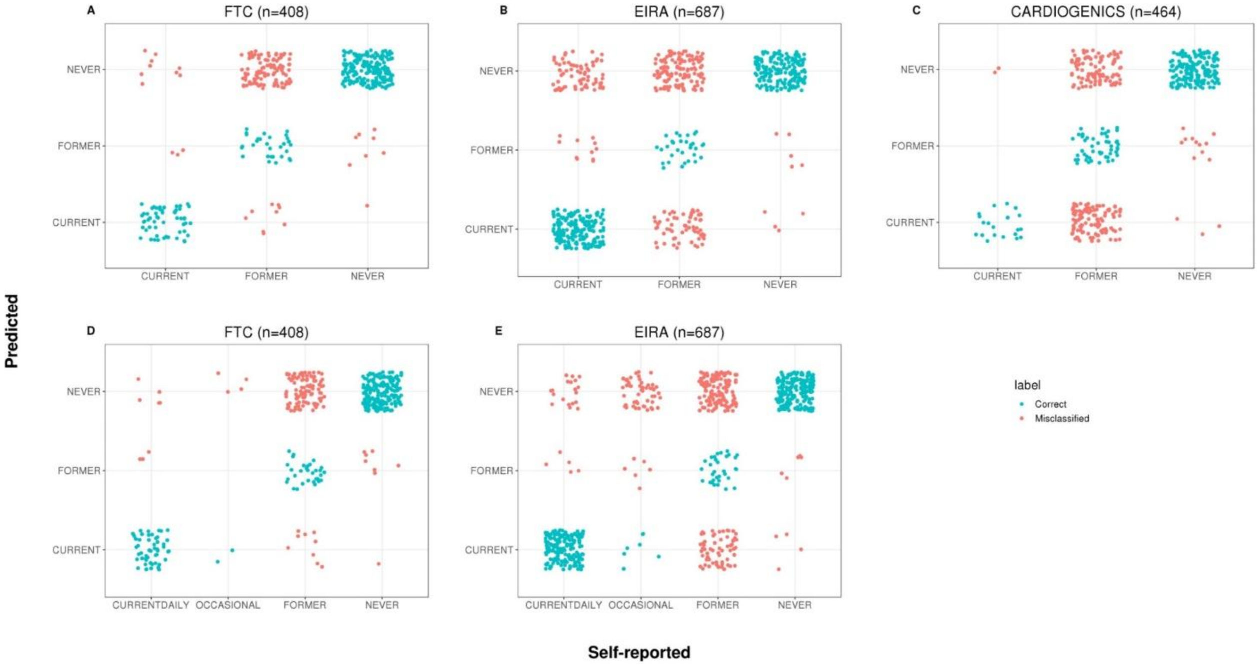

**Supplementary Figure 1: Confusion matrices comparing self-reported versus predicted smoking behavior**

Confusion matrices for FTC (**A**), EIRA (**B**) and CARDIOGENICS (**C**) datasets. Self-reported smoking behavior is compared with the predicted smoking status determined by the classifier. Points in cyan represent agreement between self-reported and predicted smoking statuses. Points in salmon represent misclassification. Based on the self-reported questionnaire data, the class of current smokers from the FTC and EIRA datasets could be further subdivided into current daily and occasional smokers. The confusion matrices **D** and **E** from FTC and EIRA show that much of the misclassification arises from the occasional smokers, who were often predicted as former or never smokers by the classifier.

**Current Vs Others**

| threshold | sensitivity | specificity |
|---|---|---|
| 0 | 0.96 | 0.46 |
| 1 | 0.94 | 0.62 |
| 2 | 0.9 | 0.74 |
| 3 | 0.9 | 0.83 |
| 4 | 0.85 | 0.89 |
| 5 | 0.82 | 0.91 |
| 6 | 0.76 | 0.94 |
| 7 | 0.72 | 0.96 |
| 8 | 0.64 | 0.97 |
| 9 | 0.54 | 0.98 |
| 10 | 0.42 | 0.99 |
| 11 | 0.33 | 0.99 |
| 12 | 0.28 | 0.99 |

| threshold | sensitivity | specificity |
|---|---|---|
| 0 | 0.82 | 0.57 |
| 1 | 0.74 | 0.67 |
| 2 | 0.71 | 0.76 |
| 3 | 0.67 | 0.81 |
| 4 | 0.65 | 0.86 |
| 5 | 0.6 | 0.9 |
| 6 | 0.55 | 0.94 |
| 7 | 0.52 | 0.96 |
| 8 | 0.47 | 0.97 |
| 9 | 0.38 | 0.97 |
| 10 | 0.29 | 0.98 |
| 11 | 0.24 | 0.99 |
| 12 | 0.18 | 0.99 |

| threshold | sensitivity | specificity |
|---|---|---|
| 0 | 0.91 | 0.6 |
| 1 | 0.91 | 0.67 |
| 2 | 0.91 | 0.7 |
| 3 | 0.86 | 0.75 |
| 4 | 0.77 | 0.78 |
| 5 | 0.73 | 0.81 |
| 6 | 0.68 | 0.85 |
| 7 | 0.59 | 0.88 |
| 8 | 0.5 | 0.9 |
| 9 | 0.45 | 0.93 |
| 10 | 0.36 | 0.95 |
| 11 | 0.27 | 0.96 |
| 12 | 0.23 | 0.97 |

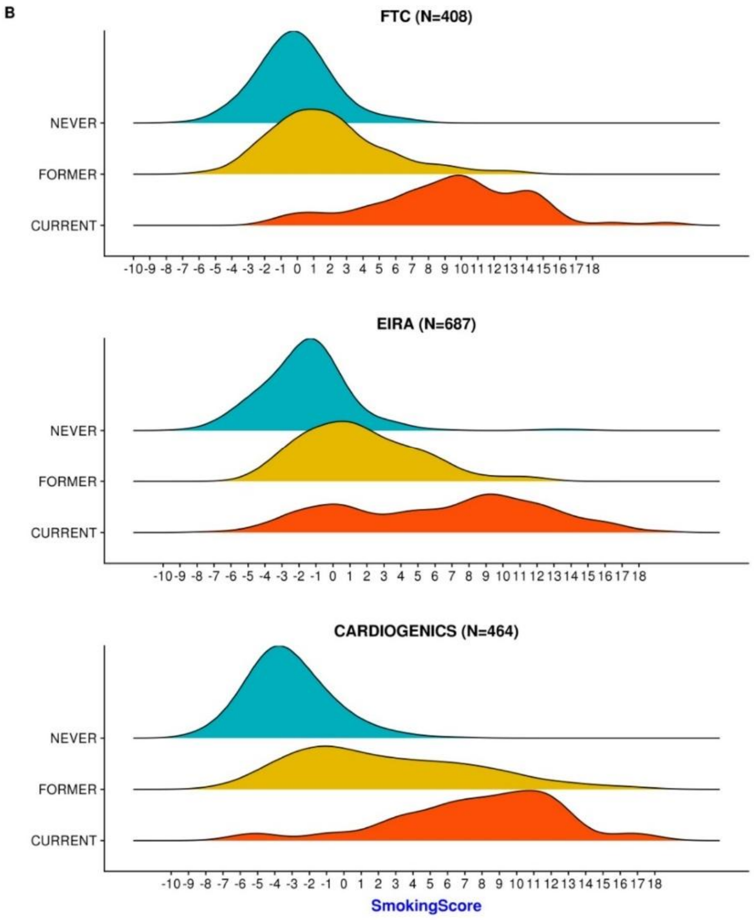

**Supplementary Figure 2: Determining an optimal threshold value for the smoking score (SSc) across the datasets**

Density plots from three test datasets with corresponding sensitivity and specificity values for different threshold values of the SSc to discriminate current vs others (former and never). A threshold value of zero seems to achieve higher sensitivity across all the datasets, however, this threshold also yields lower specificity.

**A**

**Current Vs Never**

| threshold | sensitivity | specificity |
|---|---|---|
| -5 | 0.61 | 0.98 |
| -5.5 | 0.69 | 0.97 |
| -6 | 0.7 | 0.94 |
| -6.5 | 0.79 | 0.93 |
| -7 | 0.82 | 0.89 |
| **-7.5** | **0.87** | **0.84** |
| -8 | 0.91 | 0.79 |
| -8.5 | 0.94 | 0.66 |
| -9 | 0.96 | 0.48 |
| -9.5 | 0.99 | 0.29 |
| -10 | NA | 0.13 |

| threshold | sensitivity | specificity |
|---|---|---|
| -5 | 0.55 | 0.9 |
| -5.5 | 0.6 | 0.87 |
| -6 | 0.65 | 0.82 |
| -6.5 | 0.68 | 0.76 |
| -7 | 0.73 | 0.7 |
| **-7.5** | **0.78** | **0.62** |
| -8 | 0.82 | 0.5 |
| -8.5 | 0.88 | 0.39 |
| -9 | 0.94 | 0.24 |
| -9.5 | 0.97 | 0.12 |
| -10 | 1 | 0.05 |

| threshold | sensitivity | specificity |
|---|---|---|
| -5 | 0.77 | 0.79 |
| -5.5 | 0.77 | 0.74 |
| -6 | 0.86 | 0.69 |
| -6.5 | 0.86 | 0.66 |
| -7 | 0.86 | 0.63 |
| **-7.5** | **0.91** | **0.59** |
| -8 | 0.91 | 0.5 |
| -8.5 | 0.95 | 0.37 |
| -9 | NA | 0.21 |
| -9.5 | NA | 0.1 |
| -10 | NA | 0.05 |

**B**

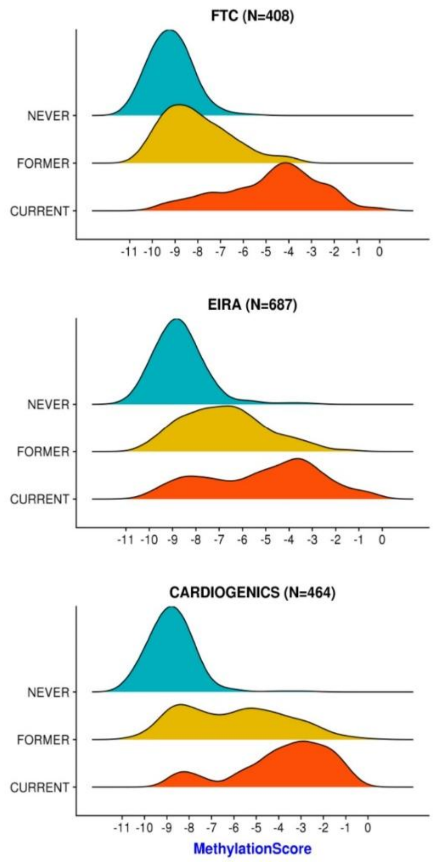

**C**

**Former Vs Never**

| threshold | sensitivity | specificity |
|---|---|---|
| -6 | 0.13 | 0.82 |
| -6.5 | 0.15 | 0.79 |
| -7 | 0.23 | 0.78 |
| -7.5 | 0.34 | 0.75 |
| -8 | 0.4 | 0.72 |
| -8.5 | 0.57 | 0.63 |
| -9 | 0.68 | 0.46 |
| -9.5 | 0.84 | 0.3 |
| -10 | 0.95 | 0.13 |

| threshold | sensitivity | specificity |
|---|---|---|
| -6 | 0.3 | 0.61 |
| -6.5 | 0.42 | 0.59 |
| -7 | 0.52 | 0.56 |
| -7.5 | 0.62 | 0.51 |
| -8 | 0.75 | 0.44 |
| -8.5 | 0.82 | 0.33 |
| -9 | 0.9 | 0.2 |
| -9.5 | 0.95 | 0.11 |
| -10 | 0.99 | 0.05 |

| threshold | sensitivity | specificity |
|---|---|---|
| -6 | 0.51 | 0.89 |
| -6.5 | 0.55 | 0.88 |
| -7 | 0.6 | 0.88 |
| -7.5 | 0.66 | 0.86 |
| -8 | 0.73 | 0.76 |
| -8.5 | 0.82 | 0.59 |
| -9 | 0.92 | 0.37 |
| -9.5 | 0.97 | 0.18 |
| -10 | 0.98 | 0.08 |

**Supplementary Figure 3: Determining an optimal threshold value for the methylation score (MS) across the datasets**

Density plots from test datasets with corresponding sensitivity and specificity values for different threshold values of the MS. **A** Different threshold values to discriminate current vs never smokers. A threshold value of -7.5 seems to achieve higher sensitivity across all the datasets. **B** Density plots with methylation score on X-axis and self-reported smoking status on Y-axis. **C** Different threshold values to discriminate former vs never smokers. It was not feasible to determine a single threshold value that can be applied to all datasets to achieve reasonable sensitivity and specificity values.

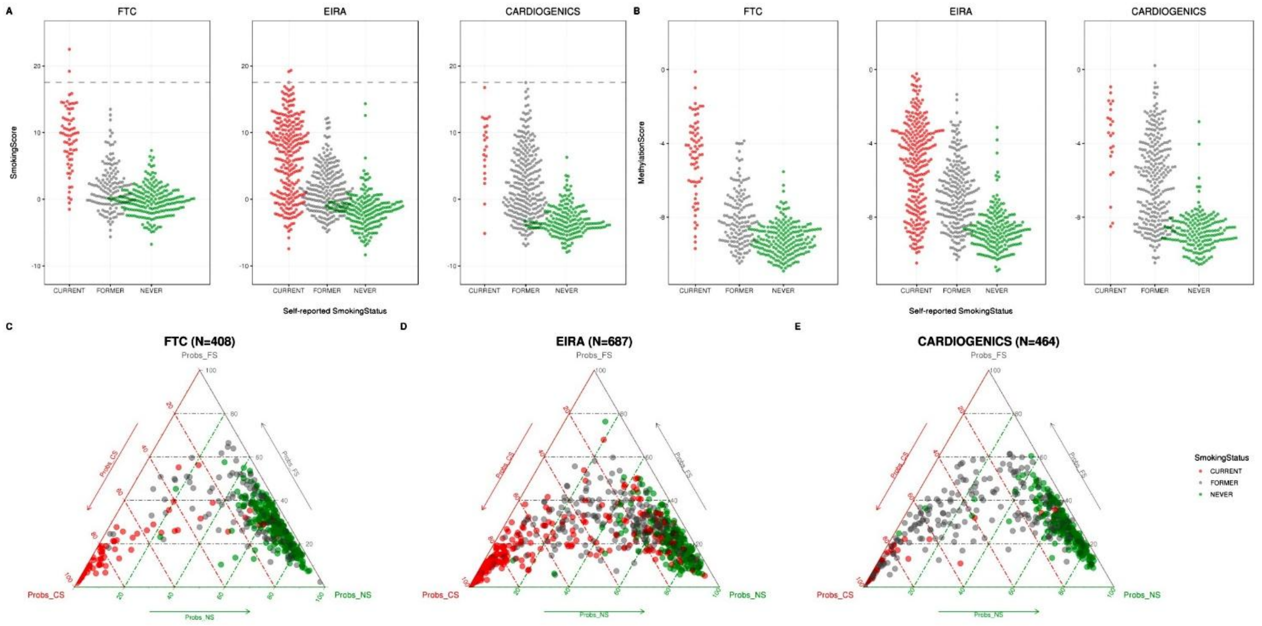

**Supplementary Figure 4: Comparison of smoking score, methylation score and smoking status classifier results across three datasets**

Bee swarm plots and ternary plots comparing SSc, MS and SSt classifier results from three datasets: FTC, EIRA and CARDIOGENICS. **A** SSc computed for current, former and never classes of the three test datasets. The grey dashed line represents the European threshold of 17.55 for current smokers proposed by Elliott and colleagues (2014). Only two current smokers from both FTC and EIRA datasets have a smoking score above 17.55. **B** MS computed for current, former and never classes of the same three datasets. Each point in the bee swarm plots corresponds to an individual, with the x-axes representing the self-reported smoking behavior category and the y-axes corresponding to the smoking or methylation score. Ternary plots **C-E** demonstrate the predicted probabilities given by our classifier in the three test datasets. Ternary plot is an equilateral triangle with each corner corresponding to a smoking status category. Each point in the ternary plot represents an individual with the corresponding triplet of probabilities that adds up to 100%. So the higher the probability for a category the closer is the point to the corresponding corner. The color of the points corresponds to self-reported smoking behavior.

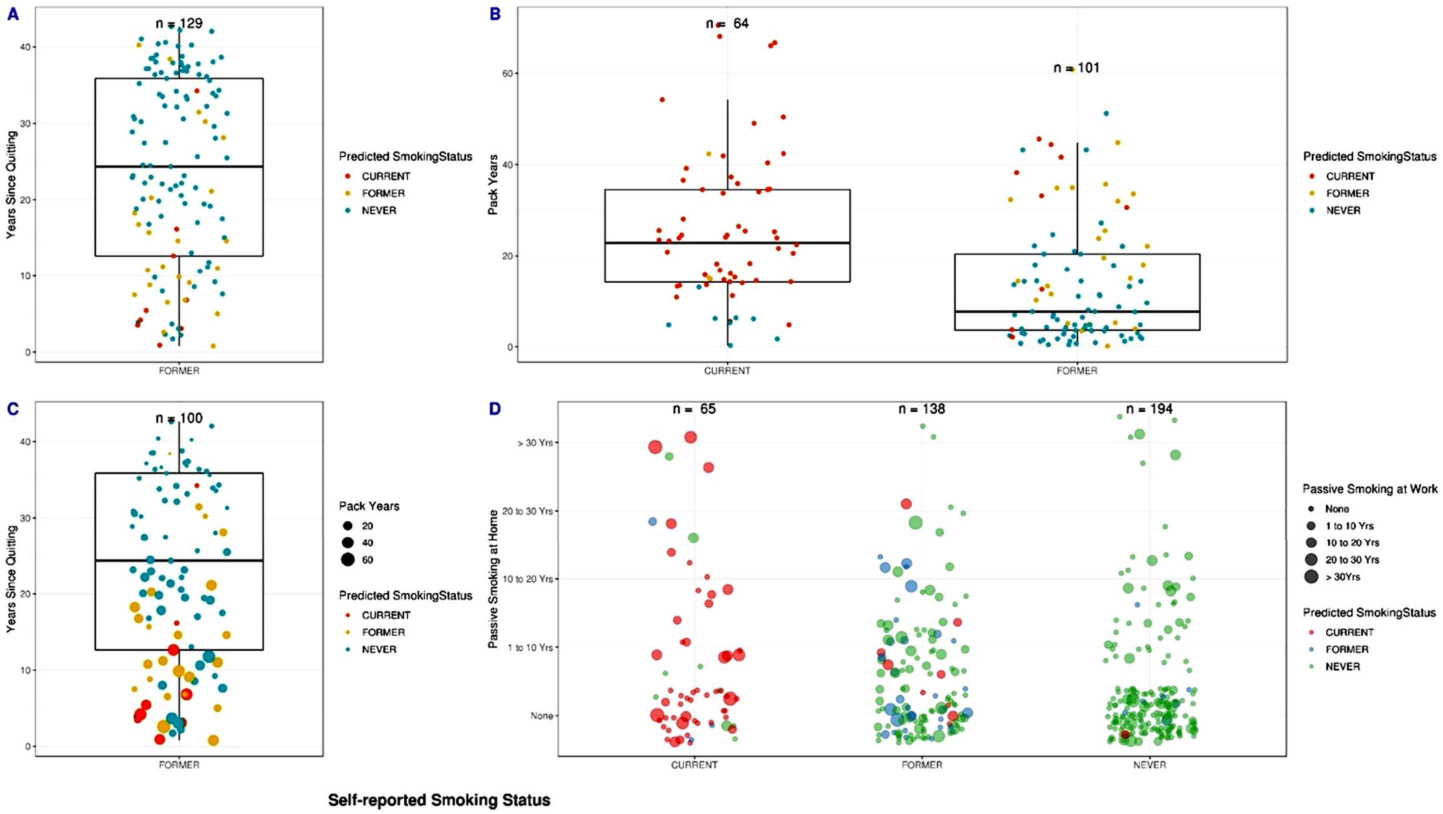

**Supplementary Figure 5: The effect of the time since abstinent and cumulative smoking exposure on the SSt classification.**

The FTC dataset (n=408) with comprehensive smoking phenotype available was used to assess the reasons for misclassification. **A** Box and whisker plot showing duration of smoking abstinence (years since quitting) of the self-reported former smokers. Here each point indicates a subject. Most of the former smokers with longer cessation time were predicted as never smokers by the classifier. **B** Box and whisker plot showing pack years from current and smokers. Here each point indicates a data point. Current and former smokers with low pack-years were predicted as never smokers by the classifier. **C** Box and whisker plot showing duration of smoking abstinence (years since quitting) of the self-reported former smokers. Here each point indicates a data point and the size of the point is proportional to the number of pack-years. Most of the former smokers with longer cessation time were predicted as never smokers by the classifier. **D** This categorical scatter plot shows passive smoking at home observed across different categories of smokers. Size of the point represents the passive smoking at work. This plot highlights the individuals who were exposed to passive smoking at home and work. The color of the point indicates the smoking status predicted by the classifier.

#### 3 Supplementary Tables

**Supplementary Table 2:** Characteristics of the training dataset.

| Training Dataset | Self-reported Smoking Status | | |
|---|---|---|---|
| DILGOM | Current Smokers | Former Smokers* | Never Smokers |
| N | 113 | 118 | 243 |
| Age (mean±SD) | 48.8±12.7 | 56.3±11.6 | 51.95±14.4 |
| Age range (years) | 25-72 | 25-74 | 25-74 |
| Sex (M/F) | 57/56 | 61/57 | 97/146 |
| Cotinine information (%) | 76 | 26 | 1 |
| Nationality | Finnish | | |
| Source | Whole blood | | |
| Platform | Infinium HumanMethylation450 BeadChip | | |

*Only former smokers who quit smoking at least a year ago were considered.

**Supplementary Table 3:** Characteristics of the test datasets used in this study to assess the performance of SSt classifier.

| Test Datasets | Dataset | N | Sex (M/F) | Self-reported Smoking Status Current | Former | Never | Nationality | Age (mean±SD) | Age range (years) |
|---|---|---|---|---|---|---|---|---|---|
| FTC (Finnish Twin Cohort ) | Available upon a reasonable request | 408 | 166/242 | 67 | 141 | 200 | Finnish | 62.2±4.3 | 32.3-69.7 |
| EIRA (Epidemio-logical Investiga-tion of Rheuma-toid Arthritis) | GSE42861 | 687 | 196/491 | 266 | 228 | 193 | Swedish | 51.9± 11.8 | 18-70 |
| CARDIOGENICS | GSE50660 | 464 | 327/137 | 22 | 263 | 179 | French and British | 55.4±6.7 | 38-67 |

**Supplementary Table 4:** Characteristics and performance measures of buccal and PBMC test datasets used to demonstrate the cross-tissue applicability of the SSt classifier.

| Source Tissue | Dataset | N | Self-reported Smoking Status | | Sex (M/F) | Age (mean) | Age range (years) | Current Vs Never | |
|---|---|---|---|---|---|---|---|---|---|
| | | | Current | Never | | | | Sensitivity (%) | Specificity (%) |
| Buccal | GSE94876 | 80[a] | 40[b] | 40[b] | 80/0 | 47.2±7.9 | 35-60 | 95 | 97 |
| PBMCs | GSE53045 | 111[c] | 50 | 61 | 0/111 | 48.4 ± 10 | NA | 89 | 96 |

[a] Of these 80 individuals, 58 were Caucasian, 21 were African-American and 1 unknown ethnicity. [b] In buccal tissue data smoking behavior was defined as cigarette smoker and non-tobacco smoker. [c] African-American ancestry.NA: Not available.

**Supplementary Table 6:** Prediction estimates from SSt classifier, across the training and test datasets.

| Datasets | Sensitivity (%) | Specificity (%) |
|---|---|---|
| Training Dataset (DILGOM) (N=474) | | |
| Current vs others | 75 | 98 |
| Former vs others | 60 | 99 |
| Never vs others | 99 | 72 |
| Test Datasets | | |
| Finnish Twin Cohort (FTC) (N=408) | | |
| Current vs others | 82 | 97 |
| Former vs others | 22 | 96 |
| Never vs others | 96 | 47 |
| EIRA (N=687) | | |
| Current vs others | 69 | 84 |
| Former vs others | 14 | 97 |
| Never vs others | 95 | 58 |
| CARDIOGENICS (N=464) | | |
| Current vs others | 91 | 73 |
| Former vs others | 19 | 95 |
| Never vs others | 92 | 65 |

Smoking status categories were predicted using the classifier, based on the 121 CpGs chosen by multinomial LASSO. Predicted smoking status was compared with self-reported smoking status. For each smoking status category, the relative sensitivity and specificity values were calculated by comparing that one with the other two categories (one vs all approach).

**Supplementary Table 7:** Methylation-based smoking status estimation approaches compared in this study.

| Outcome | Approach | Normalisation | Reference |
|---|---|---|---|
| Smoking Status (SSt) | Weighted sum of 121 CpGs, Sex and intercept terms. These CpGs were identified by Multinomial LASSO (ML). | Quantile Normalization (QN) | Our classifier |
| Smoking Score (SSc) | Weighted sum of 187 CpGs. These CpGs were identified in an EWAS (Zeilinger, et al., 2013). | Subset Quantile Normalization (SQN) | Elliott *et al* (2014) |
| Methylation Score (MS) | Weighted sum of 4 CpGs. These CpGs were selected via stepwise regression from results of an EWAS (Zhang, et al., 2016). | Illumina Normalization (ILN) | Zhang *et al* (2016) |

**Supplementary Table 8:** Prediction estimates from SSt approach when only self-reported current daily smokers were included in the current smoker category.

| Dataset | Smoking Status | Sensitivity (%) | Specificity (%) |
|---|---|---|---|
| FTC (n= 402) | Current Daily | 87 | 97 |
| | Former | 22 | 96 |
| | Never | 96 | 48 |
| EIRA (n=621) | Current Daily | 88 | 84 |
| | Former | 14 | 97 |
| | Never | 95 | 64 |

#### 4 Declarations

##### 4.1 Funding information for datasets

The DILGOM study has been supported by the Academy of Finland Research (Grants 255935 to Prof Markus Perola) for DNA methylation data. Data collection of the Finnish twin cohort samples has been supported by the Academy of Finland Center of Excellence in Complex Disease Genetics (Grants 213506 and 129680), the Academy of Finland (Grants 265240, 263278, and 312073 to JK), NIH Grant DA12854 (to Dr Pamela Madden), Sigrid Juselius Foundation (to JK), Global Research Award for Nicotine Dependence, Pfizer (to JK), the Wellcome Trust Sanger Institute, UK and the Broad Institute of MIT and Harvard, USA.

#### 4.2 Ethics approval and consent to participate

The DILGOM/FINRISK participants have provided written informed consent. The FINRISK study was designed and conducted according to the principles of the Declaration of Helsinki. Approval for the FINRISK study protocol was obtained from ethics committees of the hospital districts of Helsinki and Uusimaa, Finland.
Ethics permissions for the Finnish Twin Cohort study were obtained from the Ethics committee of the Department of Public Health, University of Helsinki and Helsinki University Central Hospital. Informed consent was obtained from all the participants. All the other datasets used in this study have obtained permissions from their respective ethical boards.

#### 4.3 Availability of data and material

Samples and data from participants of the DILGOM 2007survey are available through The National Institute for Health and Welfare's (THL) Biobank. For further information please contact the THL Biobank.
The FTC data is not publicly available due to the restrictions of informed consent. However, the FTC data is available through the Institute for Molecular Medicine Finland (FIMM) Data Access Committee for authorized researchers who have IRB/ethics approval and an institutionally approved study plan. Note that anonymized individual-level data can only be released after the study has been approved by the Research Ethics Committee of the University of Helsinki and must be carried out in collaboration with FTC investigators. To ensure the protection of privacy and compliance with national data protection legislation, a data use/transfer agreement is needed, the content and specific clauses of which will depend on the nature of the requested data. For further information please contact Jaakko Kaprio.
All the other test datasets used in this study are publicly available from the Gene Expression Omnibus (GEO).

#### 4.4 Author contributions

MO, SA and SB conceived the project. SA supervised the development of the classifier. SB implemented the analyses and designed the R package. SB and SA drafted the manuscript. MO provided critical feedback and helped shape the analysis and manuscript. SA and MO jointly supervised this work. TK contributed to the FTC and DILGOM smoking phenotypes. JK is the PI of the FTC and co-designed phenotype and DNA collection for the FTC. All authors discussed the results and commented on and approved the final manuscript.

#### Supplementary References

Aryee, M.J., *et al.* (2014) Minfi: a flexible and comprehensive Bioconductor package for the analysis of Infinium DNA methylation microarrays, *Bioinformatics*, **30**, 1363-1369.
Broms, U., *et al.* (2012) Diurnal Evening Type is Associated with Current Smoking, Nicotine Dependence and Nicotine Intake in the Population Based National FINRISK 2007 Study, *J. Addict. Res. Ther.*, **S2**.
Clarke, E. and Sherrill-Mix, S. (2017) ggbeeswarm: Categorical Scatter (Violin Point) Plots.
Cutrona, C.E., *et al.* (2005) Neighborhood context, personality, and stressful life events as predictors of depression among African American women, *J Abnorm Psychol*, **114**, 3-15.
Dogan, M.V., *et al.* (2014) The effect of smoking on DNA methylation of peripheral blood mononuclear cells from African American women, *BMC Genomics*, **15**, 151.
Elliott, H.R., *et al.* (2014) Differences in smoking associated DNA methylation patterns in South Asians and Europeans, *Clinical Epigenetics*, **6**, 1-10.
Friedman, J., Hastie, T. and Tibshirani, R. (2010) Regularization Paths for Generalized Linear Models via Coordinate Descent, *Journal of statistical software*, **33**, 1-22.
Hamilton, N.E. and Ferry, M. (2018) ggtern : Ternary Diagrams Using ggplot2, *J. Stat. Softw.*, **87**.
Huber, W., *et al.* (2015) Orchestrating high-throughput genomic analysis with Bioconductor, *Nat. Methods*, **12**, 115.
Inouye, M., *et al.* (2010) Metabonomic, transcriptomic, and genomic variation of a population cohort, *Mol. Syst. Biol.*, **6**, 441.
Inouye, M., *et al.* (2010) An immune response network associated with blood lipid levels, *PLoS Genet.*, **6**, e1001113.
Kaprio, J. (2013) The Finnish Twin Cohort Study: an update, *Twin Res. Hum. Genet.*, **16**, 157-162.
Kassambara, A. (2018) ggpubr: 'ggplot2' Based Publication Ready Plots.
Lehne, B., *et al.* (2015) A coherent approach for analysis of the Illumina HumanMethylation450 BeadChip improves data quality and performance in epigenome-wide association studies, *Genome Biol.*, **16**, 37.
Liu, Y., *et al.* (2013) Epigenome-wide association data implicate DNA methylation as an intermediary of genetic risk in rheumatoid arthritis, *Nat. Biotechnol.*, **31**, 142-147.
Morris, T.J., *et al.* (2014) ChAMP: 450k chip analysis methylation pipeline, *Bioinformatics*, **30**.
Prasad, G.L., *et al.* (2016) A cross-sectional study of biomarkers of exposure and effect in smokers and moist snuff consumers, *Clin. Chem. Lab. Med.*, **54**, 633-642.

Stolt, P., *et al.* (2003) Quantification of the influence of cigarette smoking on rheumatoid arthritis: results from a population based case-control study, using incident cases, *Ann. Rheum. Dis.*, **62**, 835-841.
Touleimat, N. and Tost, J. (2012) Complete pipeline for Infinium(®) Human Methylation 450K BeadChip data processing using subset quantile normalization for accurate DNA methylation estimation, *Epigenomics*, **4**, 325-341.
Tsaprouni, L.G., *et al.* (2014) Cigarette smoking reduces DNA methylation levels at multiple genomic loci but the effect is partially reversible upon cessation, *Epigenetics*, **9**, 1382-1396.
Wickham, H. (2016) *ggplot2: Elegant Graphics for Data Analysis*. Springer.
Wilke, C.O. (2018) cowplot: Streamlined Plot Theme and Plot Annotations for 'ggplot2'.
Zeilinger, S., *et al.* (2013) Tobacco smoking leads to extensive genome-wide changes in DNA methylation, *PLoS One*, **8**.
Zhang, Y., *et al.* (2016) Self-reported smoking, serum cotinine, and blood DNA methylation, *Environ. Res.*, **146**, 395-403.
